# Sex and Alternative Splicing in Disease: a meta-analytic approach to identify interactions

**DOI:** 10.64898/2026.05.29.728908

**Authors:** Netanya Keil, Alison Morse, Colin Callahan, Patrick Concannon, Lauren M McIntyre

## Abstract

How cell type, sex and disease interact and affect gene expression and splicing is an important, but complicated question. Visualizing and testing specific hypotheses around these complex interactions is an important first step to identifying molecular components underpinning complex disease. Using a meta-analytical framework, we develop an analytical path for identifying testable molecular hypotheses of complex interactions between splicing, sex, disease and cell type. We focus on type 1 diabetes (T1D) but the approach is generalizable to any complex disease with defined candidate loci. Previous studies report T1D-associated splicing in candidate genes, differences in disease effects across immune cell types, sex effects on splicing and cell-type-specific splicing. However, identifying and interpreting complex interactions between sex, splicing and disease are challenging. Here we demonstrate how a gene expression study of T1D, designed to evaluate these interactions can be analyzed in a straightforward manner. We find that sex-dependent T1D-associated splicing is markedly more prevalent in CD4⁺ T cells than in CD8⁺ T cells, affecting 72% of T1D candidate genes in CD4⁺ cells compared to 30% in CD8⁺ cells. We pinpoint exons whose rate of inclusion is affected by the interaction of sex and disease. We use long-read RNAseq to identify novel intron retention events and splice sites which are quantified with short-reads leading to a richer description of the regulatory impact of T1D on alternative splicing. We identify a set of candidate isoforms for follow-up molecular studies in *BACH2*, a transcription factor known to be relevant in disease prevalence.

## Main Text

Alternative splicing, a mechanism that regulates transcriptome and proteome diversity, is a molecular phenotype associated with several genetically complex diseases including cancer ^1^, neurodegenerative disease^2^, autoimmune disease generally^3; 4^ and type 1 diabetes in particular.^5^ Alternative splicing is cell type and tissue specific^6–8^ and there is evidence for sex-specific alternative splicing in some contexts.^9–11^ Evaluating the potential interactions between cell type, sex, and disease in alternative splicing is currently stymied by the lack of good tools for relevant biological interpretations of these multi-factor interactions.

Autoimmune diseases are a group of genetically complex disorders characterized by loss of immune tolerance leading to impairment or destruction of host organs or tissues. Many autoimmune disorders exhibit a significantly higher prevalence in females as compared to males.^12; 13^However for some autoimmune disorders, notably Type 1 Diabetes (T1D), evidence for a sex bias in prevalence is mixed. ^14–16^ T1D results from autoimmune destruction of the Beta cells of the pancreas resulting in complete dependence on exogenously administered insulin for survival. Phenotypes of the disorder, such as rate of disease progression from initial evidence of autoimmunity to frank diabetes, or the prevalence of complications and comorbidities do demonstrate an effect of sex ^17; 18^ Females have also been shown to have a higher rate of all-cause mortality when compared to males.^19^ There are credible biological models that suggest that sex effects contribute to T1D disease.^20^ However, because of the focus on prevalence as a primary determinant of risk factors, sex has often been omitted in analyses of other molecular phenotypes for T1D.^21^

T1D has distinct cell-type specific manifestations. Cell transfer studies in mouse models of T1D suggest that both CD4+ and CD8+ T cells are necessary for the autoimmune destruction of pancreatic beta cells.^22; 23^ Cytotoxic T cells (CD 8+) are key effectors of beta cell destruction as they are able to recognize autoantigens on Beta cells and directly destroy them through the secretion of cytotoxic molecules and pro-inflammatory cytokines and the initiation of apoptosis pathways.^24^ Helper T cells (CD4+) mediate beta cell destruction by recognizing islet autoantigens and secreting cytokines.^25^ CD4+ and CD8+ T cells have also been shown to exhibit different gene expression and splicing profiles in mice.^8^

Linkage and genome-wide association studies (GWAS) have identified more than 100 chromosomal regions that contribute significantly to T1D risk.^26; 27^ In studies of splicing quantitative trait loci (sQTL), T1D risk variants are associated with alternative splicing in T cells.^5^ Mechanistic studies of individual T1D risk variants have identified alleles that regulate specific splicing events associated with a molecular phenotype that may contribute to disease. For example, an allele at SNP rs1893592 is associated with retention of intron 9 in the *UBASH3A* gene, resulting in elevated IL2 production in CD4+ T cells.^28; 29^ A credible causative variant in a T1D risk locus at chromosome 20p13, rs6043409, alters a coding region splicing enhancer altering the distribution of spliced SIRPG isoforms and consequently the protein expression levels of the cell surface receptor CD172G.^30^ In addition, two SNPs in the T1D risk gene *PTPN22* have been shown to affect splicing of the *PTPN22* transcripts.^31–33^ In addition to these examples of splicing effects associated with T1D genetic risk loci, there is also accumulating evidence for genetic effects on splicing in pancreatic Beta cells that may contribute to T1D risk.^34; 35^ However, whether sex interacts with T1D disease associated splicing has not been examined.

### In this study, we use a combination of third generation long read sequencing and short read RNA Seq of CD4+ and CD8+ T cells derived from T1D cases and controls to investigate the impact of sex and disease status on splicing patterns in CD4+ and CD8+ immune cells

We use long reads to catalog the expressed exons and introns in both cell types and short reads to quantify these features.^36^ Unsurprisingly CD4+ and CD8+ cells, differ in their splicing patterns^8^, and consistent with other reports using long reads we identify many more variable features than are currently annotated.^37^ We couple a logical series of hierarchical hypotheses tests with meta-analytic approaches providing intuitive visualizations that enable us to unpack the interactions between tissue, sex, splicing and disease. We find that disease affects splicing in a sex specific manner for 70% of the T1D candidate genes in CD4+ cells and 30% of the T1D candidate genes in CD8+ cells and we are able to pinpoint alternative splicing patterns for molecular follow up. Our approach is easily transferred to other disease contexts as we leverage biological structure with meta-analysis, a mature analytical framework, for hypothesis tests and visualizations of interactions between sex, disease and transcript structure.

The Type 1 Diabetes Genetics Consortium (T1DGC) ascertained families with one or more affected offspring and obtained biospecimens from parents and both affected and unaffected offspring.^38^ Subjects for the current study were selected from T1DGC resources to be unrelated with self-reported European ancestry and anonymized.^39^ European ancestry was confirmed by genotyping.^27^ Viably frozen peripheral blood mononuclear cells (PBMCs) from study subjects were heated at 37°C until nearly thawed, washed twice in washing buffer (RPMI media supplemented with 20% heat-inactivated fetal bovine serum) and incubated overnight (up to 16 hours) in 10 ml of fresh washing buffer at 37°C and 5% CO2 in a 25 cm^2^ flask. Cells were transferred to a centrifuge tube, incubated with DNase I (STEMCELL Technologies), centrifuged and resuspended in MACS buffer (Miltenyi Biotec) at 4°C. CD4^+^ and CD8^+^ T and CD19^+^ cell populations were purified by sequential positive selection on magnetic beads on an autoMACS Cell Separator (Miltenyi Biotec) according to the manufacturer’s instructions. RNA for long read sequencing was extracted using RNeasy (Qiagen). A single sample from each of the case/control male/female CD4/CD8/CD19 groups was selected at random and these 8 independent samples were used for long read sequencing. Long read libraries were prepared following the Iso-Seq Express workflow provided by PacBio using the NEBNext Single Cell/Low Input cDNA Synthesis and Amplification Module (New England BioLabs) in conjunction with the Iso-Seq Express oligo kit to generate full-length, barcoded cDNA that was pooled and shipped to the Genomics and Cell Characterization Core Facility at the University of Oregon for SMRT Bell library preparation and sequencing.

We followed the Isoseq3 long-read pre-processing pipeline. Circular consensus reads (CCS) (pbccs/5.0.0) were demultiplexed and primers were removed using Lima (v2.7.1). CCS reads had on average 30 subreads. Reads without one full pass were not considered further. PolyA tails and concatemers were removed using Isoseq refine (v.3.4.0 https://isoseq.how/ ). Bam files were converted to fastq (bedtools v2.29.2) and mapped to the Ensembl hg38 reference (release 104) with minimap2 (v.2.24 -ax splice -C5 –secondary=no). CCS reads were combined across samples and clustered using Isoseq3 cluster to create Isoseq transcript models (v.3.4.0 https://isoseq.how/). Isoseq transcript models were mapped to the Ensembl hg38 reference (release 104) with minimap2 (v.2.24 -ax splice -C5 –secondary=no). We evaluated the Isoseq transcript models and the CCS reads with SQANTI3QC and transcript models inconsistent with the CCS reads were removed.^40^ The remaining full-length transcripts consistent with either Refseq (p13) or Ensembl (release 104) annotations were used to identify the expressed exons in the gene model and variable features (alternative donors/acceptors and introns). Features at least 10 nucleotides long were annotated in a bed file (https://data.rc.ufl.edu/pub/mcintyre/t1d_metaanalysis/isoseq_transcripts/).^36; 41^

Short read sequencing libraries were prepared using the NEBNext Ultra II Directional RNA Library Prep kit (New England BioLabs) and sequenced by Macrogen. There were very few successful CD19+ selections, and therefore this cell type was not considered further. The number of CD4+ and CD8+ samples included in each group is provided in Table S1. The 2x150 paired-end Illumina short reads were trimmed using Cutadapt (v2.1), and R1 and R2 reads were merged if they overlapped using BBMerge (bbmap v38.44). Unmerged reads were aligned paired-end, and merged reads were aligned single-end to the hg38 genome (release 104) using BWA mem (v0.7.17). PCR duplicate reads were removed using Samblaster (v 0.1.24), and reads with unique primary alignments were quantified using a bed file of feature annotations to calculate the average coverage per nucleotide (APN) for each feature.^36; 41^ A feature was considered detected for a given cell-type/sex/case status if the average coverage per nucleotide (APN) was greater than 5 for 50% or more of the samples. Features not detected in at least one of the four conditions (Male Case, Male Control, Female Case, Female Control) in a cell type (CD4^+^ or CD8^+^) were not considered further.

For each sample the Q1, mean, median and Q3 for the feature APN was calculated. Samples were scaled to adjust for differences in coverage using the upper quartile normalization.^42^ For features detected in the control samples for both cell types we fit the model: (Y_ijk_)=m+Age_i_+Sex_j_+CellType_k_ +sex*Celltype_jk_+error_ijk_ where Y_ijk_ is the log2 of the APN for individual i with sex (j=male,female) and cell type k (k=CD4^+^, CD8^+^) and tested the null hypothesis that the expression of feature f was equal for males and females in cell type k using a contrast. For features detected in either cell type we fit the model Y_ijk_=m+Age_i_+Sex_j_+CaseStatus_l_ +sex*CaseStatus_jl_+error_ijk_ where Y_ijl_ is the log2 of the APN for individual i with sex (j=male,female) and CaseStatus l (l=diabetic, control). We evaluated statistical significance using a nominal significance level of 0.05 as our goal was to minimize false negatives to evaluate broad trends between cell types and T1D status, and not to make inferences about individual features *per-se* (https://data.rc.ufl.edu/pub/mcintyre/t1d_metaanalysis/model_output/).

To evaluate whether T1D has an effect on splicing, we computed an effect size for each feature f in each cell type k in males (ES_male_fk_) and females (ES_female_fk_) separately as 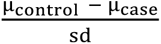 (. Under the null hypothesis of no T1D effect on splicing, the effect sizes should be consistent across features. In contrast, if T1D alters splicing in males/females, we expect heterogeneity in the T1D effect, reflecting changes in relative exon/intron inclusion that depends on the case status. We tested for heterogeneity of feature-level effect sizes for each sex using Cochran’s Q test for heterogeneity, as implemented in the metafor^43^ fixed-effect meta-analysis framework in each cell type separately. We test the null hypothesis of equal effects of disease on splicing between the sexes, by testing whether the estimated difference in effect sizes between sexes (ES_male_ij_ - ES_female_ij_) was consistent across all features for each cell type using Cochran’s Q test for heterogeneity with the estimation of variance as the weighted average between the sexes.

Long-read RNA sequencing (LR-RNASeq) exposes the limitations of existing annotations.^41; 44^ From the observed long read transcripts (t=110,579), 51% of them included a novel 5’ and 3’ splice junctions and/or evidence of intron retention. For example, in T1D candidate gene *UBASH3A* we observed intron retentions spanning exons 3-6 of the annotated MANE transcript (Figure 1B). In UBASH3A we also observed several alternate 5’ and 3’ splice sites in exons 7, 9 and 11 of the annotated MANE transcript (Figure 1B). By defining features from long reads we capture features of the transcripts expressed in our cohort that would have otherwise been missed.

**Figure 1:**
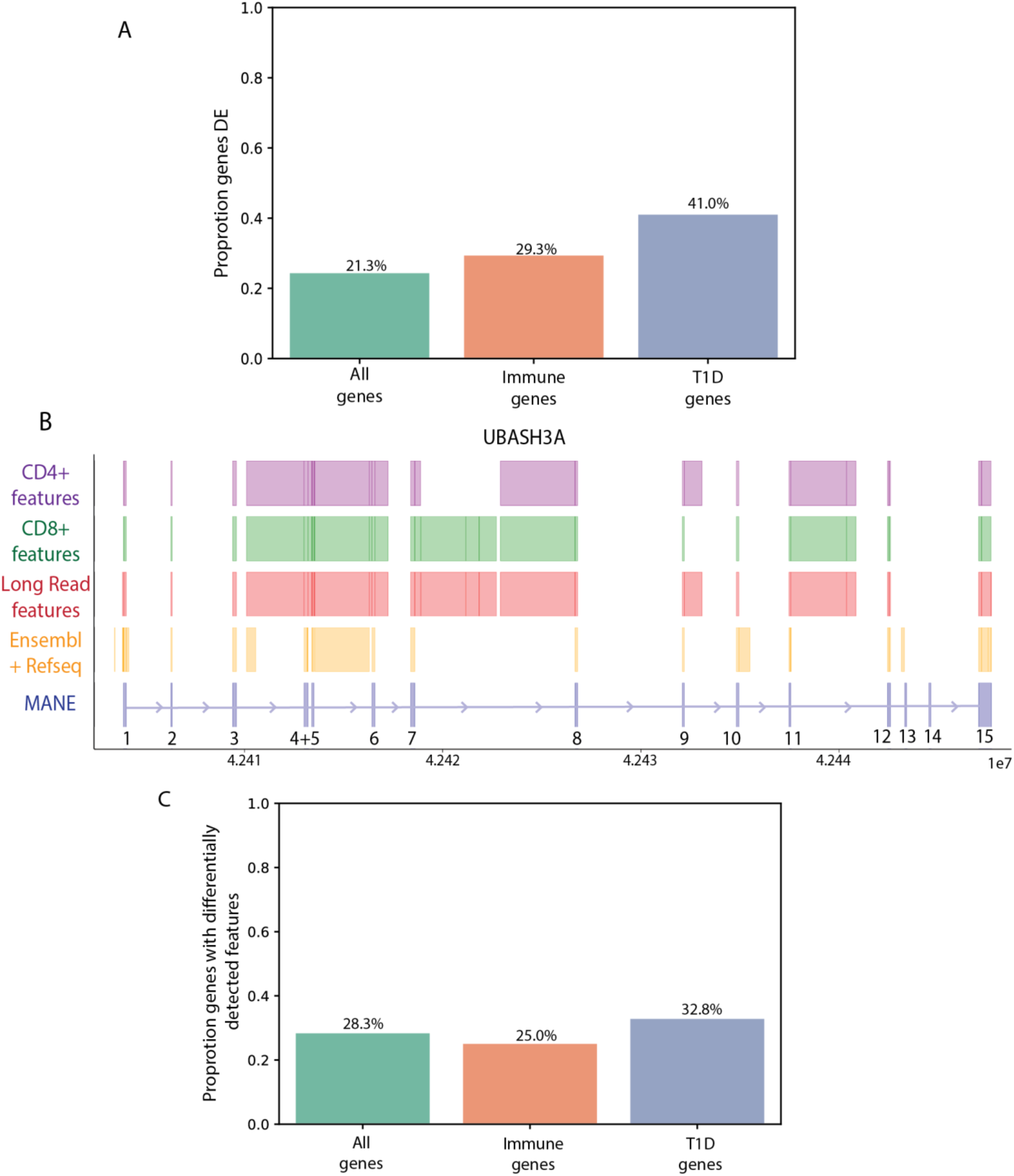
CD4+ and CD8+ T cells have different transcriptomic profiles. Gene expression in control samples was compared between CD4+ and CD8+ samples (A) Proportion of all genes, immune genes and T1D genes that are differentially expressed between CD4+ and CD8+ control samples. T1D candidate genes and immune relevant genes are more likely to be differentially expressed between these cell types. (B) Annotated and detected exon and intron features for T1D candidate gene UBASH3A. From bottom to top i) The reference MANE transcript ii) all annotated features in Refseq/Ensembl iii) Exon/Intron features detected by long read sequencing in both cell types iii) Exon/Intron features detected in CD4+ RNA-seq samples (n=113) iv) Exon/Intron features detected in CD8+ RNA-seq samples (n=98). Exon Region 5 of UBASH3A is differentially detected between CD4+ and CD8+. (C) Proportion of all genes, immune genes and T1D genes with at least one differentially detected feature between CD4+ and CD8+samples. T1D genes are more likely to have differentially detected exon/intron features between CD4+ and CD8+ cells.

Do CD4+ and CD8+ T cells exhibit distinct splicing profiles in normal controls? CD4+ and CD8+ T cells both contribute to T1D pathogenesis, through distinct immunological functions. CD4+ T cells are thought to play important roles in coordinating and sustaining autoimmune responses, whereas CD8+ T cells are more directly implicated in the destruction of pancreatic beta cells.^24;25^ Consistent with these functional differences, prior long-read analyses in mouse T cells have shown that CD4+ and CD8+ populations differ in isoform usage.^8^ In this study, differences in detection of individual features between CD4+ and CD8+ samples reflect differences in isoform usage between these cell types. In T1D-associated genes, 33% of T1D associated genes had at least one feature differentially detected between CD4+ and CD8+ compared to 30% of immune genes and 28% of genes overall. (Figure 1C) Interestingly, we observed alternative 5’ splice sites in exon 7 and 9 of UBASH3A, which are detectable in CD4+ samples but not CD8+ samples suggesting differential splice site usage between these cell types. (Figure 1A)

Is gene expression different in normal controls for CD4+ and CD8+ cells? Focusing on the features that were detected in both cell types we observed 24% of all expressed genes being differentially expressed between CD4+ and CD8+ T cells with 29% of immune-related genes and 41% of T1D-associated genes differing in expression, suggesting that the genes most relevant to immune function and T1D pathogenesis are particularly cell-type specific in their expression.

Does disease status affect splicing in T1D associated genes? The *a priori* hypothesis behind this study was that the molecular phenotype of T1D associated risk loci would differ in their molecular phenotype between cases and controls and that these molecular differences would help elucidate disease phenotype. This gene set is also a manageable size for targeted evaluation of complex interactions. We detected 60 T1D-associated genes in both cell types, and an additional gene (IL-10) in CD4+. The observed differences in expression and detection of features between CD4+ and CD8+ cells, support the stratification of the remainder of the analyses by cell type. Using the empirically identified features from the long reads, we tested for differential expression of individual features between T1D cases and controls in CD4+ and CD8+ cells. Newman et al.^45^, showed that T1D-associated splicing changes in regulatory T cells are dominated by IR events. We found both exonic and intronic features were differentially expressed at higher rates in CD4+ compared to CD8+ cells. In long reads, introns were detected in 19 of the 61 T1D candidate genes (31%) and of these 19 genes, 4 T1D genes in CD4+ cells and 1 T1D gene in CD8+ cells had a differentially expressed intron between cases and controls. With respect to exonic features, 38 T1D genes in CD4+ cells and 22 T1D genes in CD8+ cells had a differentially expressed exon. Together, these results demonstrate that T1D-associated genes exhibit case-control differences at the level of individual features, which is consistent with changes in isoform usage in individuals with T1D.

How can we test whether the pattern of disease associated splicing differs between males/females? We stratify the splicing analyses by both cell type and sex. Although feature-level analyses provide important insight into discrete splicing events, including intron retention, and exon skipping, they capture individual local differences rather than the overall pattern of splicing variation. We use the structure of the gene and a meta-analytic framework to evaluate the pattern of exon inclusion/intron retention across a gene. We treat each feature as a ‘study’ and sort the features based on the gene model from the 5’ to 3’ end. We calculate the expression level and compute sex specific effect sizes for each feature (Figure 2 B-D). Effect sizes enable the visualization and evaluation of changes in the magnitude and direction of the effect of T1D between sexes. This is illustrated by the T1D candidate gene *RPAP2* in CD4+ cells, where the effect of T1D differs both across the gene and by sex (Figure 2 E-G). In ER1 (the first exonic region), T1D has opposite effects in males and females. In ER4 (the fourth exonic region), a T1D effect is evident in females but not in males. In ER7 (the seventh exonic region), T1D affects both sexes in the same direction, but the magnitude of the effect differs (Figure 2 E-G). The impact of T1D varies across individual features in a sex-dependent manner. To capture this variation quantitatively we use Cochran’s Q test to test for heterogeneity of effect sizes. In RPAP2, both males and females are significant for the test of heterogeneity, indicating that both males and females exhibit differential splicing between T1D cases and controls (Figure S1).

**Figure 2:**
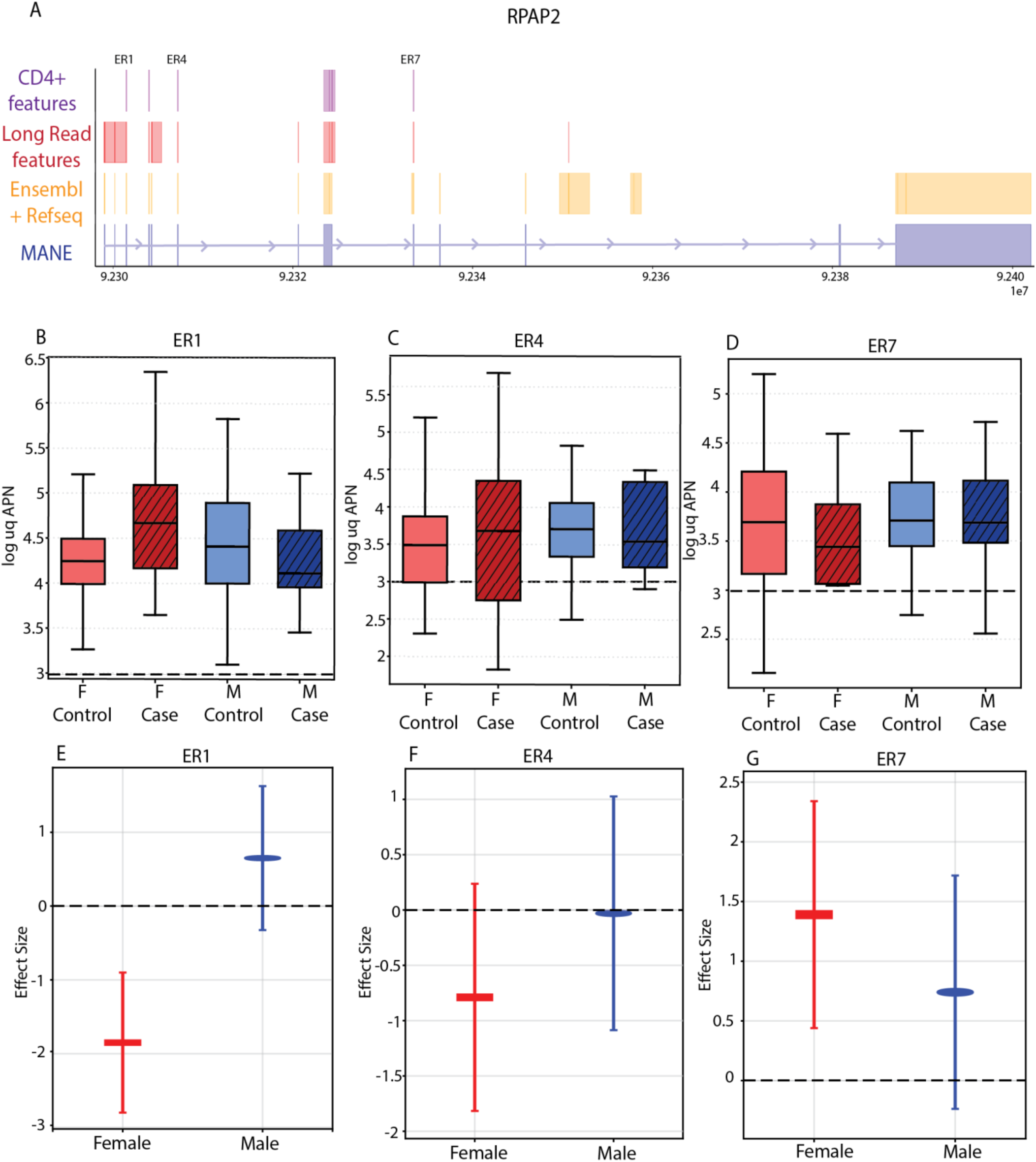
Effect of T1D is exon dependent. A) Annotated and detected exon and intron patterns for T1D candidate gene RPAP2. From bottom to top i) The reference MANE transcript ii) all annotated features in either Refseq/Ensembl iii) Exon/Intron features detected by long read sequencing in both cell types iv) Exon/Intron features detected in CD4+ RNA-seq samples (n=113) Exon regions 1,4 and 7 are labelled. (B) – (D) Normalized expression level (log uq APN) of exon regions 1,4 and 7 stratified by sex and T1D status. Log UQ APN of 3 is shown with black dotted line across all plots. (E)-(G) Effect size of T1D on exon regions 1,4 and 7 stratified by sex in CD4+ RNA-seq samples. A positive effect size indicates that controls are more highly expressed than cases and a negative effect size indicates that cases are more highly expressed than controls for a given feature. Effect size of 0 is represented by the black dotted line. The upper and lower bounds of the 95% confidence interval for the effect size are shown for each feature.

How can we evaluate the potential underlying three-way interaction between sex, splicing and T1D? In STRN4 (Figure 3B), we observed significant heterogeneity of T1D effect sizes across exons in females but not males in CD4+ samples, indicating a female-specific effect of T1D on splicing for this gene (Figure 3B). This pattern where a T1D relevant gene is differentially spliced in cases versus controls in females but not in males is observed in 17 T1D genes in CD4+ cells and 4 T1D genes in CD8 (Figure S2 A&B). Conversely, we observed 2 T1D genes in CD4+ cells and 4 T1D genes in CD8+ cells with evidence of differential splicing in cases versus controls in males but not in females (Figure S2A&B). In the *STRN4* gene, there is a three-way interaction in CD4+ and CD8+ cells (Figure S3). In CD4+ cells, two features (ER1:EF1, ER7:EF1) have pronounced effect size differences between males and females in CD4+ samples (Figure 3B, Figure S3) and are likely driving the observed significant interaction effect. At the feature level both ER1:EF1 and EF7:EF1 have a significant sex*T1D effect (Table S2). An overwhelming majority (72%) of the T1D candidate genes in CD4⁺ T cells and 30% of the T1D-associated genes in CD8⁺ T cells show evidence for a T1D × sex × splicing interaction (Figure 4A).

**Figure 3:**
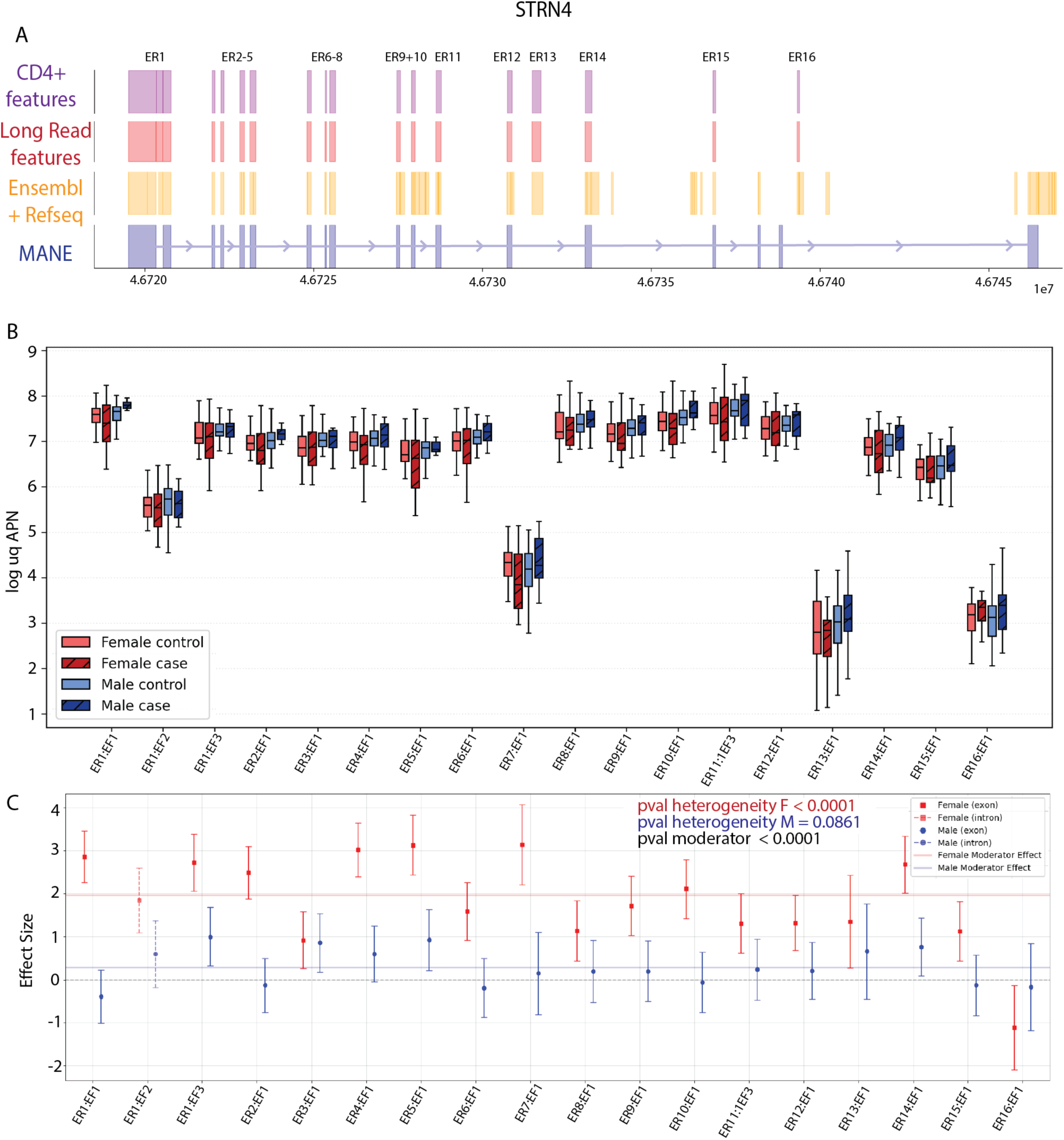
Effect of T1D on splicing is sex dependent. (A) Annotated and detected exon and intron patterns for T1D candidate gene STRN4. From bottom to top i)The reference MANE transcript ii) all annotated features in either Refseq/Ensembl iii) Exon/Intron features detected by long read sequencing in both cell types iv) Exon/Intron features detected in CD4+ RNA-seq samples (n=113) (B) Normalized expression level (log_uq_apn) of CD4+ RNASeq samples for all features of the T1D candidate gene STRN4. There are differential rates of exon and intron feature inclusion across the length of the transcript (C) Effect size of T1D on all exon/intron features in STRN4 stratified by sex in CD4+ RNA-seq samples. The upper and lower bounds of the 95% confidence interval for the effect size are shown for each feature. Pvalue for the test of heterogeneity of female and male effect sizes and the Pvalue for the test of a moderator (sex) effect on the whole gene is shown. The test for heterogeneity is significant for females but not for males indicating that there is differential splicing between T1D cases and controls in female samples for the STRN4 gene. The test for moderator (sex) effect is significant indicating that there is an overall effect of sex on the gene.

**Figure 4:**
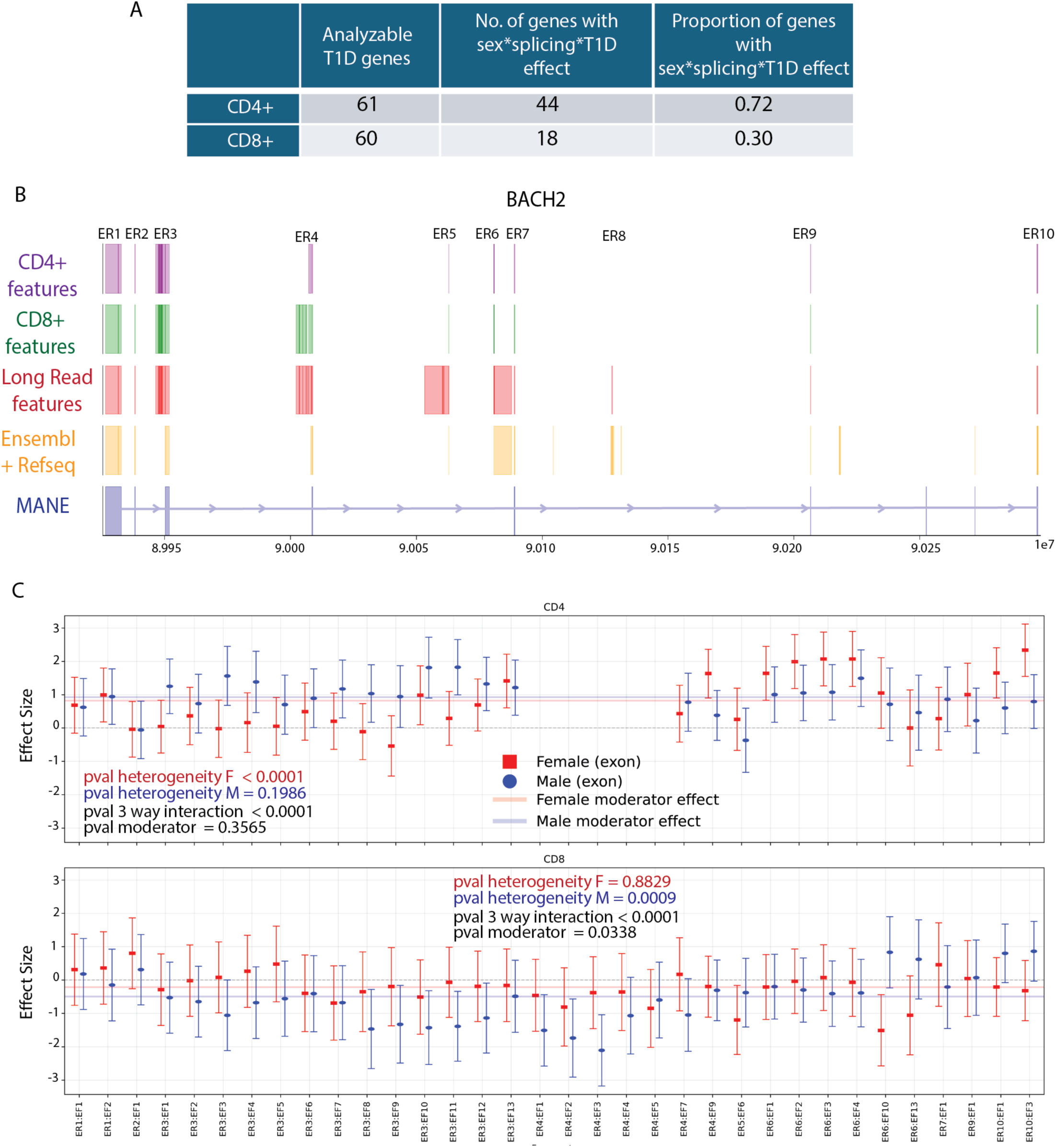
Effect of T1D on splicing is cell type dependent. A) Number and proportion of analyzable T1D genes with a 3-way interaction effect interaction of sex,splicing and T1D status for CD4+ and CD8+ samples. A higher proportion of T1D genes have a sex*splicing*T1D effect in CD4+ samples compared to CD8+ samples B) Annotated and detected exon patterns for T1D candidate gene BACH2. From bottom to top i) The reference MANE transcript ii) all annotated exons in Refseq/Ensembl iii) Exon features detected by long read sequencing in both cell types iii) Exon features detected in CD4+ RNA-seq samples (n=113) iv) Exon feautures detected in CD8+ RNA-seq samples (n=98). C) Effect size of T1D on all exonic features in BACH2 stratified by sex in CD4+ RNA-seq samples (top) and CD8+ RNA-seq samples (bottom). The upper and lower bounds of the 95% confidence interval for the effect size are shown for each feature. Pvalue for the test of heterogeneity of female and male effect sizes and the Pvalue for test of a moderator (sex) effect on the whole gene is shown. In CD4, the test for heterogeneity is significant for females but not for males indicating that there is differential splicing between T1D cases and controls in female samples for the BACH2 gene. The inverse effect is observed in CD8+ samples where we observe an effect of T1D on splicing in males but not females. Pvalue for the test of the 3-way interaction of sex, T1D and splicing differences is shown. This test of heterogeneity tests for the interaction of sex and splicing. Both CD4+ and CD8+ have a 3-way T1D*sex*splicing effect in the BACH2 gene.

How do we interpret these cell type specific interaction effects in a biological context? The T1D candidate gene *BACH2* illustrates the framework for such interpretations. Exon region 3 is differentially detected between CD4⁺ and CD8⁺ samples, consistent with baseline differences in isoform usage between these cell types (Figure 4B). Both CD4⁺ and CD8⁺ cells exhibit a T1D × sex × splicing interaction for *BACH2* (Figure S4), but the nature of the interaction differs by cell type. In CD4+ cells, differential splicing between T1D cases and controls is observed in females but not males, whereas in CD8⁺ cells the inverse occurs, with differential splicing present in males but not females (Figure 4C). Together, these findings show that CD4⁺ and CD8⁺ T cells differ substantially in how sex modifies the splicing response to T1D, highlighting cell-type-specific regulatory mechanisms underlying disease related splicing changes and point to specific regions of the transcript for further examination and molecular follow-up.

In this study, we used a combination of long and short read RNA sequencing to characterize splicing in T1D-associated genes in CD4+ and CD8+ T cells and to evaluate how splicing is altered by the interaction between T1D disease status and sex. By empirically defining features from the long-read data, rather than relying on reference annotation, we focus on features present in these cell types including retained introns and alternative splicing donor and acceptors not present in the reference annotation. This enabled us to evaluate variable features between CD4+ and CD8+ cells, T1D cases and controls and sexes. For example, in the gene *BACH2* gene we observe alternative detection of splice sites between cell types, and we observe alternate usage of splice sites based on sex and case status (Figure 4 B&C).

Despite arising from the same lymphoid lineage, CD4+ and CD8+ T cells display clear differences in both gene expression and splicing in normal controls. These differences are particularly frequent in genes previously associated with T1D risk suggesting that elements of disease phenotype may be associated with these molecular phenotypes. More genes showed the three-way interaction- sex*splicing*T1D in CD4+ cells than in CD8+ cells demonstrating that CD4+ and CD8+ cells represent distinct molecular contexts for disease. One interpretation is that CD4+ cells are more transcriptionally responsive to the regulatory consequences of T1D, consistent with their central roles in immune coordination and adaptive response.^46^

Complex interactions including the three-way interactions between sex, disease, and splicing are inherently difficult to interpret. Our visualizations are built on the molecular structure of each gene, with observed exons and introns providing the framework for both visualization and statistical testing. By layering a heterogeneity-based statistical approach onto the biological structure, we can quantitatively evaluate whether the effect of T1D is consistent across the gene. The visualizations borrowed from the meta-analytic framework make it possible to identify specific exons or introns where the effect of T1D status differs from other features, and to determine whether variation in the effect of T1D differs by sex. The corresponding heterogeneity test complements these visualizations by providing a gene-level statistical framework for identifying genes in which the effect of T1D is not constant across features, consistent with differential splicing.

Meta-analysis provides an intuitive way to visually interpret complex interaction patterns at the level of specific features, complemented by feature level significance testing. By linking affected exons to annotations of functional domains and/or locations of risk variants it is possible to generate mechanistic hypotheses for follow-up molecular studies. A limitation of this approach is that it is best suited to a targeted or limited set of genes, where feature-level visualization and interpretation can be carried out in a biologically meaningful way.

Sex is as an important modifier of T1D-associated splicing, indicating that the molecular response to T1D differs between the sexes and sex needs to be evaluated in future studies. Prior work has reported differences between males and females in incidence, age of onset^18^, comorbidities, and long-term outcomes^19; 47^, with some studies showing that women with T1D experience a disproportionately elevated risk of vascular and cardiovascular complications.^17; 48;49^ Sex-specific immune differences have also been reported at the molecular level, including differences in circulating cytokine, chemokine and growth factor profiles among individuals with T1D.^50^ Although our study was not designed to test the clinical consequences of alternative splicing directly, our results raise the possibility that differential splicing of T1D-relevant genes contributes to these sex-dependent differences in disease presentation or progression. More broadly, they support the idea that sex may shape not only the epidemiology of T1D, but also the molecular programs through which T cells respond to disease.

An additional consideration is what stage of disease these transcriptomic differences represent. The T1D samples were collected after diagnosis, therefore the splicing patterns we observe could reflect established disease, a residual signature of earlier immune dysregulation, or both. Therefore, these changes may mark processes involved in disease initiation, progression or chronic adaptation. This question is especially relevant given that only a small fraction of circulating T cells are likely to be directly engaged in active islet autoimmunity at any one time. Regardless, these are reproducible differences between cases and controls suggesting that the splicing changes identified here are not confined to rare autoreactive CD4+ cells but instead may reflect a broader immune response to T1D or a persistent remodeling of T cell state in the context of disease.

We provide a practical and easily deployable meta-analytic framework for identifying and interpreting disease-associated splicing using feature-level visualization and gene-level heterogeneity testing. Although applied here to T1D-associated genes, the underlying biological premise is general: genes are composed of a set of ordered individual exon and intron features, and context specific splicing will result in changes in the relative use of these features that can be detected by testing for heterogeneity. Our framework can be applied broadly to other diseases, cell types, or experimental settings in which alternative splicing is suspected to contribute to phenotype. In this way, the approach provides an intermediate layer between large-scale gene discovery approaches and time and resource intensive targeted molecular studies. By pinpointing individual exons and introns with the differential response across biological groups, molecular hypotheses can be developed. Because meaningful interpretation requires detailed gene-by-gene evaluation of feature-level patterns this approach is best suited to a limited or targeted set of genes.

## Data and code availability

Raw sequencing data generated and analyzed in this study are not publicly available due to participant privacy concerns and restrictions related to human subject data. Processed feature-level count matrices and analysis outputs are available at https://data.rc.ufl.edu/pub/mcintyre/t1d_metaanalysis/. The code used in this study is available at https://github.com/McIntyre-Lab/papers/tree/master/keil_t1d_meta_analysis_2026.

## Acknowledgements

We thank the patients and families of the Type 1 Diabetes Genetics Consortium (T1DGC) for their participation and contribution to this study. We thank Anna Yang for assistance with generating the visualizations used in this manuscript. We also thank Dr. Jeremy Newman for initial quality control and processing of the long-read and short-read RNA-seq data. This work was supported by R01DK116954 awarded to P.C. from the National Institute of Diabetes and Digestive and Kidney Diseases (NIDDK, https://www.niddk.nih.gov/).

## Author contributions

Conceptualization: PC, LM; Data curation: NK, AM; Formal Analysis: NK, AM, LM; Funding acquisition: PC, LM; Investigation: NK, AM, CC; Methodology: NK, AM, LM; Project administration: PC, LM; Resources: ; Software: NK, AM; Supervision: LM; Visualization: NK; Writing – original draft: NK, LM; Writing – review & editing: NK, PC, LM

## Declaration of interests

The authors declare no competing interests.

## Web resources

metafor R package; https://cran.r-project.org/web/packages/metafor/

## Notes

### Competing Interest Statement

The authors have declared no competing interest.

