## Supplemental Tables and Figures for "Sex and Alternative Splicing in Disease: a meta-analytic approach to identify interactions"

**Table S1.** Short-read RNA-seq sample counts by cell type, sex and T1D case status

| **Cell Type** | **Sex** | **Case status** | **Count** |
| --- | --- | --- | --- |
| CD4 | Male | Control | 46 |
| CD4 | Male | Case | 13 |
| CD4 | Female | Control | 39 |
| CD4 | Female | Case | 15 |
| CD8 | Male | Control | 38 |
| CD8 | Male | Case | 13 |
| CD8 | Female | Control | 35 |
| CD8 | Female | Case | 12 |

| featureID | Pvalue sex*T1D interaction |
| --- | --- |
| ENSG00000090372:ER10:EF1 | 0.111603329 |
| ENSG00000090372:ER11:EF1 | 0.434452327 |
| ENSG00000090372:ER12:EF1 | 0.417956908 |
| ENSG00000090372:ER13:EF1 | 0.62983266 |
| ENSG00000090372:ER14:EF1 | 0.168309746 |
| ENSG00000090372:ER15:EF1 | 0.351405043 |
| ENSG00000090372:ER16:EF1 | 0.491619197 |
| ENSG00000090372:ER1:EF1 | 0.017341066 |
| ENSG00000090372:ER1:EF2 | 0.369053984 |
| ENSG00000090372:ER1:EF3 | 0.223319991 |
| ENSG00000090372:ER2:EF1 | 0.055182814 |
| ENSG00000090372:ER3:EF1 | 0.988808454 |
| ENSG00000090372:ER4:EF1 | 0.080640377 |
| ENSG00000090372:ER5:EF1 | 0.115192218 |
| ENSG00000090372:ER6:EF1 | 0.187503692 |
| ENSG00000090372:ER7:EF1 | 0.029569738 |
| ENSG00000090372:ER8:EF1 | 0.493092332 |
| ENSG00000090372:ER9:EF1 | 0.268507955 |

**Table S2.** Pvalue for the test of interaction of sex and T1D status for all features in the STRN4 gene (Ensembl gene ID ENSG00000090372). ER1:EF1 and ER7:EF1 are highlighted in yellow

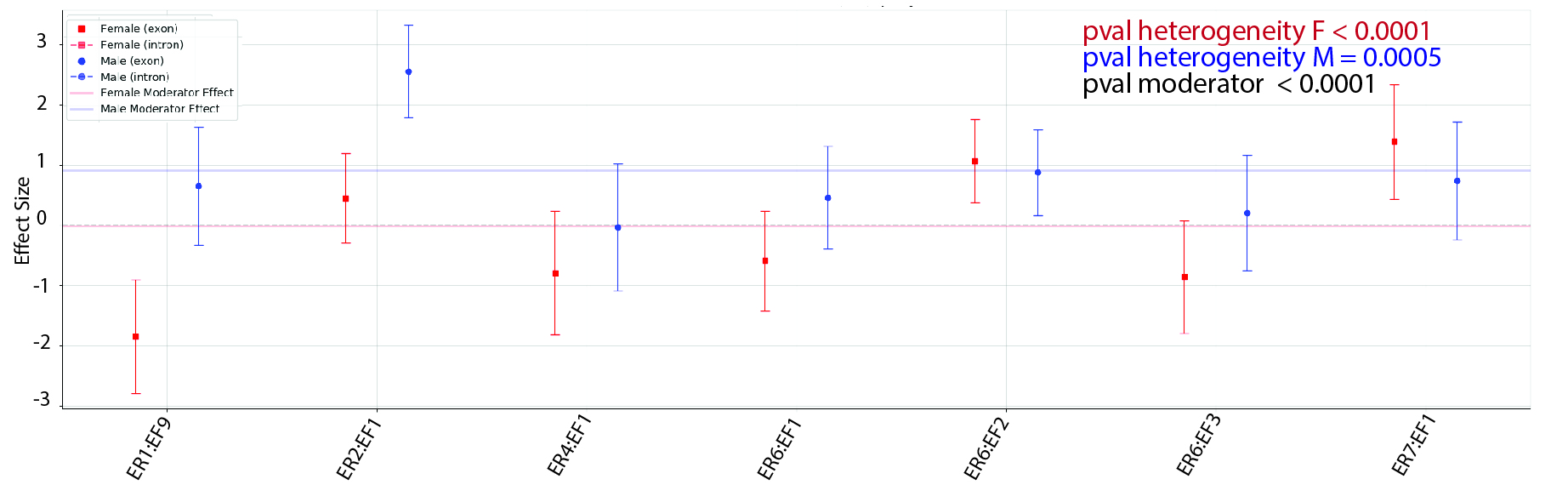
**Figure S1:** Effect size of T1D on all exon regions and segments in RPAP2 stratified by sex in CD4+ RNA-seq samples. Pvalue for the test of heterogeneity of female and male effect sizes and the Pvalue for test of a moderator (sex) effect on the whole gene is shown. The test for heterogeneity is significant for both males and females indicating that there is differential splicing between T1D cases and controls in both sexes. The test for sex (moderator) effect is significant indicating that there is an overall effect of sex on the gene.

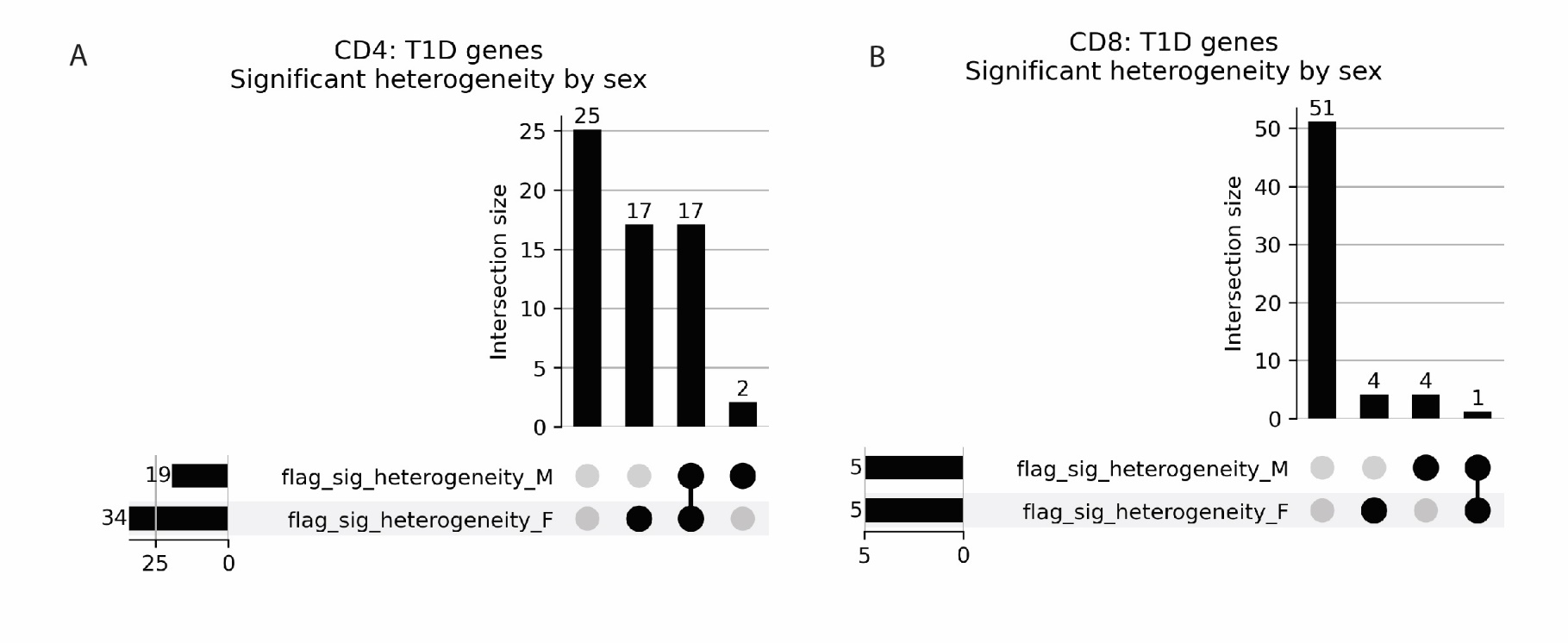

**Figure S2**: Sex-specific heterogeneity of T1D-associated genes in CD4+ and CD8+ T cells. Upset plots show the overlap of T1D candidate genes with significant feature-level heterogeneity in males and females for CD4+ samples (A) and CD8+ samples(B). Black connected dots indicate the sex-specific heterogeneity category represented by each bar, and bar height indicates the number of genes in each intersection. In CD4+ samples, significant heterogeneity was observed in both males and females for 17 genes, with 2 genes significant only in males and 17 genes significant only in females. In CD8+ T cells, we observed 4 genes significant only in females, and 4 genes significant only in females and 1 gene significant in both sexes.

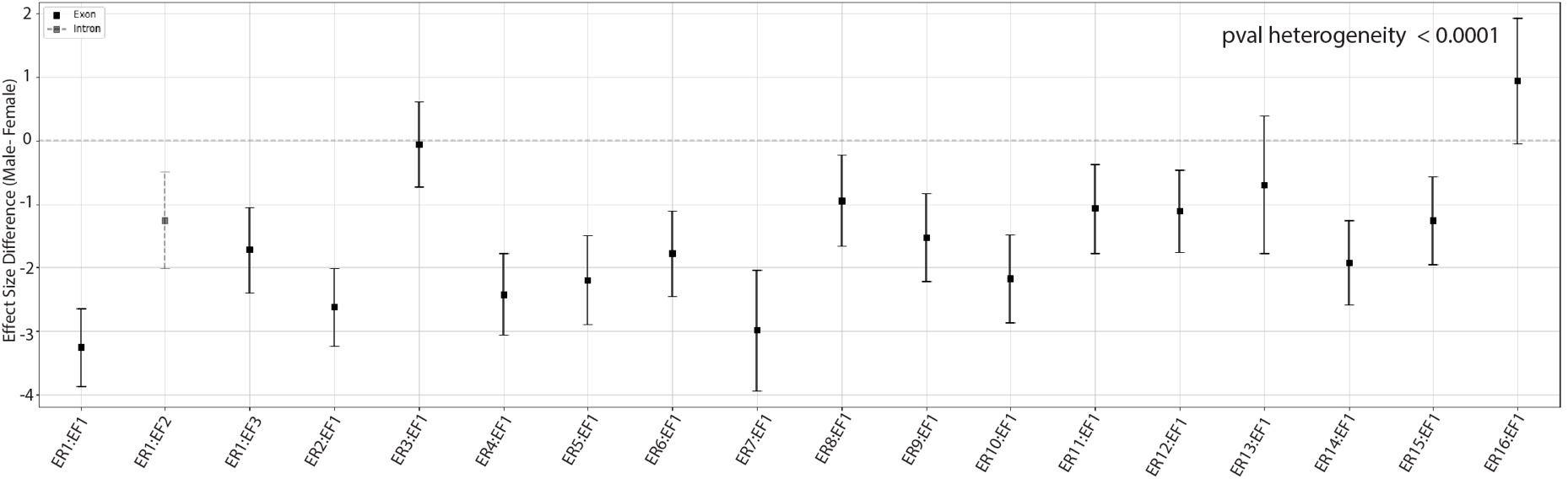
**Figure S3**: Difference of effect size (male ES – female ES) of T1D for all exonic and intronic features in STRN4 in CD4+ RNA-seq samples. Pvalue for the test of heterogeneity of effect size differences is shown. Error bars indicate 95% confidence intervals based on the estimated sampling variance for each feature. This test of heterogeneity tests for the three-way interaction of sex,splicing and T1D status. STRN4 is significant for the 3 way interaction of sex, T1D and splicing ind CD4+

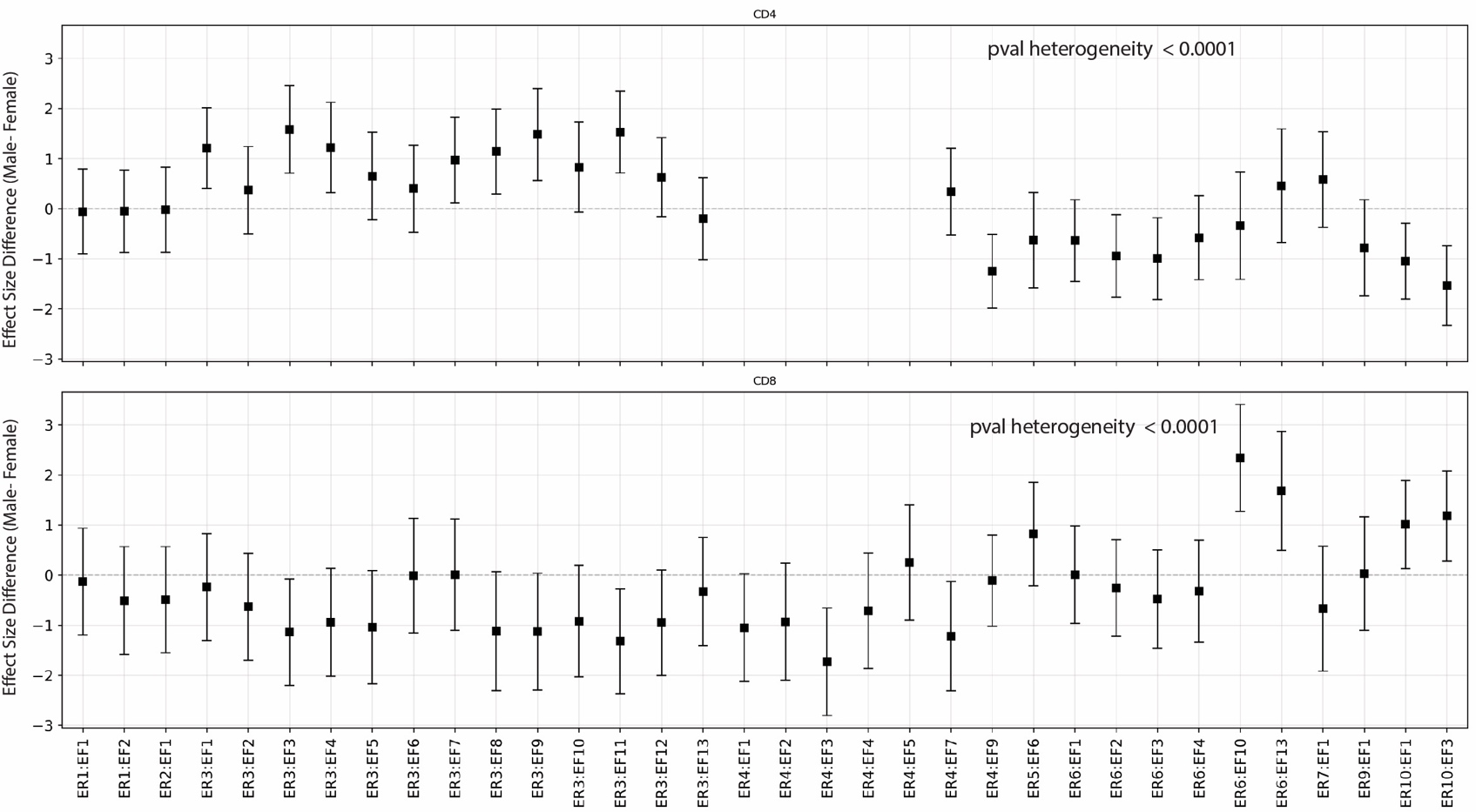

**Figure S4:** Difference of effect size (male ES – female ES) of T1D for all exonic and ifeatures in BACH2 in CD4+ (top) and CD8+ (bottom) RNA-seq samples. Pvalue for the test of heterogeneity of effect size differences is shown. Error bars indicate 95% confidence intervals based on the estimated sampling variance for each feature. This test of heterogeneity tests for the three-way interaction of sex, splicing and T1D status. BACH2 is significant for the 3 way interaction of sex, T1D and splicing in both CD4+ and CD8+ samples
